# Record-breaking wildfire intensity and nocturnal spread during the extreme 2025 wildfire season in Southwestern Europe: causes and impacts

**DOI:** 10.64898/2026.06.15.732327

**Authors:** Víctor Resco de Dios, Àngel Cunill Camprubí, Simon J Schütze, Fernando Castedo-Dorado, Juan Picos, Joaquín Ramírez, Rut Domènech, Mercedes Bachfischer, Marc Castellnou Ribau, Adrián Cardil

## Abstract

Southwestern Europe faced an extreme wildfire season in 2025, with nearly 700,000 hectares burned in the Iberian Peninsula (IP) alone. Here, we analyze the drivers and impacts of the 2025 wildfire season in the IP and its significance within the ongoing global pyrocrisis. Decades-long declines in burned area, driven by increased fire suppression, ceased after an inflection point in 2022. Fire intensity has escalated over the last two decades, and the energy emitted in 2025 approached that produced annually by a 1,000MW nuclear reactor. Despite a historically wet spring, an extreme summer heatwave triggered a flash drought, dehydrating fuels below critical thresholds. Remarkably, 29-42% of all wildfires spread faster at night than during the day, a seldom-reported phenomenon likely arising from interactions between surface weather, atmospheric instability, and pyroconvective processes. Global change-induced increases in fire intensity facilitated the overwhelming of suppression efforts during simultaneous fire events that may have been manageable decades ago. Fire activity expanded into previously fire-free high-altitude regions, and there was a marked change in fire-size distributions, with the largest wildfire in record and the largest proportion of burned area by megafires (those burning over 5,000ha). Impacts included over 2,000 premature deaths from smoke exposure and significant effects on protected areas. These results indicate shifts in key components of anthropogenic fire regimes, including unprecedented nocturnal fire acceleration and increased burned area and fire intensity, with escalating impacts on human health and ecosystems.

## Introduction

The 2025 wildfire season ranks second in burned area in the Iberian Peninsula (IP) since records began in 1980, with nearly 700,000 hectares burned, exceeded only by 2017. Current evidence identifies rural abandonment, resulting in increased fuel continuity and load, as the primary driver of any potential worsening of fire regimes, with climate change contributing through enhanced fuel aridity and fire weather (Moreira et al., 2019; Senande-Rivera et al., 2025). However, whether long-term fire activity in the IP is shifting, and the mechanisms underlying the exceptional 2025 season and its associated impacts, remain unclear.

Analyses of fire regime shifts are often conducted retrospectively, once changes have become established (Pausas and Fernández-Muñoz, 2011). However, emerging evidence suggests that key components of fire regimes may already be shifting in many regions. Globally, the size and simultaneity of the most extreme wildfires are increasing (Schütze and Resco De Dios, 2025). Some studies suggest that fire intensity has also increased, although direct quantitative evidence remains limited (Schütze and Resco De Dios, 2025). Changes in fire behavior, including increased nighttime fire intensity, have also been reported (Balch et al., 2022; Luo et al., 2024), but their role as drivers of extreme wildfire seasons remains to be explored.

Fire activity in 2025 was highly concentrated in a two-week window in early August, when most extreme wildfires co-occurred. This extreme synchronization suggests that fire simultaneity, alone or interacting with higher fire intensity, was a key driver of the season’s severity. Under this interpretation, the 2025 season was driven primarily by an increase in the number of simultaneous ignitions, straining suppression capacity during a concurrent, record-breaking heatwave (AEMET, 2025). In turn, suppression teams would have prioritized protecting populated areas, allowing fires in less-inhabited regions to spread and subsequently threaten additional communities. Fire simultaneity is increasingly recognized as a major operational challenge (Beltrán-Marcos et al., 2026; Rodrigues et al., 2019), and the synchronicity of fire activity is expected to intensify under continued warming (Jolly et al., 2015; Yin et al., 2026). Meanwhile, investment in suppression resources has risen sharply (Urbieta et al., 2019), and it remains unclear whether such a substantial increase in suppression can offset these compounding operational pressures. Evaluating whether fire simultaneity functionally overwhelmed suppression teams in 2025 and quantifying its impact relative to extreme weather and fuel connectivity remain to be tested.

Here, we test whether a shift in key components of the fire regime contributed to the extreme fire season of 2025, and assess its broader relevance in the context of global fire activity trends. Specifically, we seek to: (i) characterize the 2025 wildfire season in the Iberian Peninsula and test for significant trends of burned area, fire size, and fire intensity; ii) understand the underlying contributions of weather, fire simultaneity, and arson; iii) quantify changes in nocturnal wildfire spread rates; and (iv) evaluate associated impacts, particularly on public health and nature protection. Previous analyses of extreme fire seasons have primarily focused on aggregate burned area or on individual extreme wildfire events(Jain et al., 2024; Rodrigues et al., 2023a). Here, we integrate regional-scale patterns with the dynamics of the largest wildfires to provide a comprehensive assessment of extreme fire activity in the context of an emerging global pyrocrisis and potential shifts in fire regime components. Our results showed an unrecognized importance of nighttime fire spread.

## Methods

### Changes in burned area, ignitions and intensity

We combined different data sources of burned area. First, we used the official fire statistics for Spain (the Spanish General Fire Statistics, EGIF, https://servicio.mapa.gob.es/incendios/Search/Publico) and Portugal (from the Nature and Forests Conservation Institute, https://www.icnf.pt/florestas/gfr/gfrgestaoinformacao/dfciinformacaocartografica) since 1980, the year they started in Portugal, as this provides the longest record on burned area. We used official statistics for long-term analyses of annual area burned (Fig. 1a) and for the relationships between the number of ignitions and burned area (Fig. 5). However, these datasets are not spatially explicit and were therefore limited to non-spatial analyses. Spatial analyses of burned areas were based on data from the European Forest Fire Information System (EFFIS dataset) for the 2001–2025 period (San-Miguel-Ayanz et al., 2012). This represents an updated and harmonized data source at the subcontinental scale, based on the <u>MODIS</u> Collection 6 (C6) MCD64A1 burned area product (Giglio et al., 2018). We retrieved daily burned area data over the full period, aggregated weekly using the pyroregions often used for the IP (Rodrigues et al., 2023a), and second-level NUTS (nomenclature of territorial units for statistics) from the GADM v.4.1 dataset (https://gadm.org/). When a fire crossed multiple regions, it was assigned to the region where most of the burned area occurred. We used the Global 1-arc-second (∼30m) Digital Terrain Model (DTM) (GEDTM30) (Ho et al., 2025) to understand changes in wildfire with altitude.

**Figure 1.**
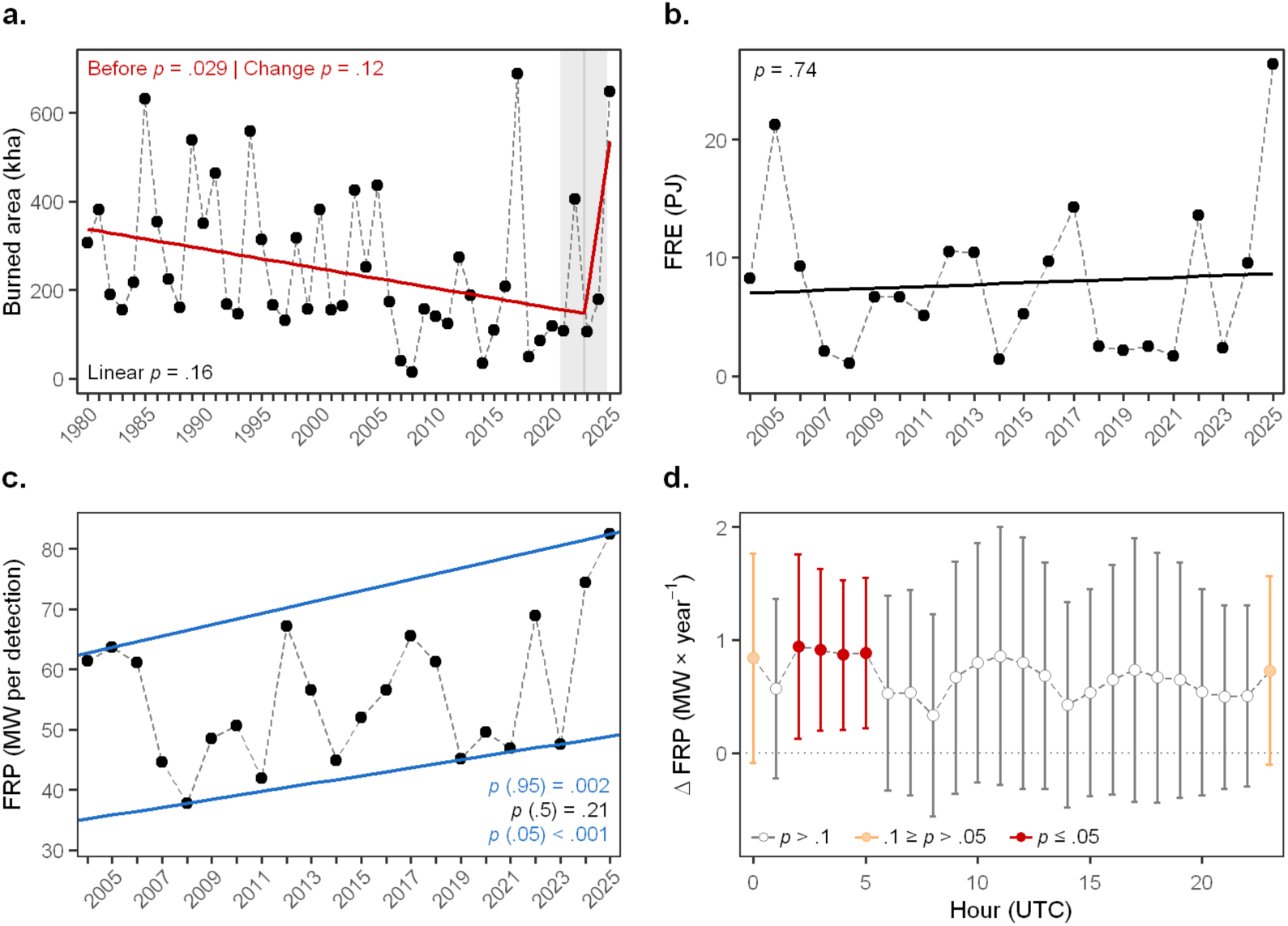
| Changes in burned area and fire intensity signal a potential change in fire regime components. The panels show annual patterns in (a) burned area; (b) fire radiative energy (FRE); and (c) mean Fire Radiative Power (FRP), a proxy for fire intensity (calculated as FRE normalized by the number of active fire detections). Panel (d) shows changes in the slope of the annual increment of mean FRP (ΔFRP) as a function of hour. In (a-c), *P*-values in black, red, and blue indicate the results of linear, segmented, and quantile (0.05 and 0.95 quantiles) regressions, respectively. Error bars in (d) represent SE, and p-values indicate significance levels in the change in slope.

We assessed changes in total fire intensity, and also in nighttime intensity from Fire Radiative Power (FRP) in the Meteosat Second Generation (MSG) Spinning Enhanced Visible and Infrared Imager (SEVIRI) (Mota and Wooster, 2018). MSG SEVIRI shows a nadir spatial sampling resolution of around 3 km and captures data every 15 minutes. We used the Fire Radiative Energy Emission (FREM) dataset built on the MSG SEVIRI sensor and available at the EUMETSAT LSA-SAF webpage (https://lsa-saf.eumetsat.int/), which provides total energy emitted. From this data, we estimated annual FRE by integrating all hourly FRE values. Mean FRP, an indicator of fire intensity, was calculated as the ratio of FRE to the number of detections, which is also available in the FREM data. We divided the FRE-detection ratio by 3600 seconds to convert energy (MJ) to power (MW).

Ignition data was collated from multiple sources, including previously mentioned official statistics whenever possible, and newspaper reports in the remaining cases.

Analyses on burned area trends, on altitudinal changes, and fire radiative energy included linear regression, Sen’s slope, segmented regression, and quantile regression, depending on whether we wanted to understand general patterns (linear regression or Sen’s slope), identify break points (segmented regression), or trends in extreme values (0.05 and 0.95 quantile regression). To test for a significant change in nighttime intensity, we calculated the hourly ΔFRP mean from the slope of the regression between the hourly FRP mean and years.

### Changes in fire rate of spread

In addition to fire intensity, we calculated and modeled fire spread rates. Calculating rates of spread requires the prior development of fire isochrones, which is notably difficult because it relies on a combination of remotely sensed data. To address this complexity, we used two approaches to reconstruct fire spread. First, we used data from the Visible Infrared Imaging Radiometer Suite (VIIRS) on board the joint NASA/NOAA Suomi National Polar-orbiting Partnership (Suomi NPP) satellite, and also used the NOAA-20 Joint Polar Satellite System 1 and the NOAA-21 Joint Polar Satellite System 2, as described in detail elsewhere (Cardil et al., 2023b). The VIIRS 375 m active fire product uses a multi-spectral algorithm to identify fire activity across 5 imagery channels (I-bands), 16 moderate resolution channels (M-bands), and the Day/Night Band (DNB). The second, complementary approach further uses Fire Radiative Power data from the Flexible Combined Imager (FCI) aboard the Meteosat Third Generation (MTG), with a spatial sampling resolution of around 1 km and data captured every 10 minutes (Xu et al., 2026).

The rate of spread using VIIRS data was calculated as in previous publications (Cardil et al., 2023b). That is, we clustered VIIRS detections into fire perimeter isochrones and then derived ROS as the ratio of the length of the longest axis (in km) of the ellipse fitted to the burned pixels to the fire duration (in hours). Thanks to the multiple satellites from VIIRS, we could obtain 2 passes per day within 1–2 hours around 01:30h and 13:30h. To characterize differences between nighttime and daytime ROS, we compared ROS calculated during nighttime overpasses and compared it against that calculated using daytime overpasses. However, this data only captures a snapshot of fire spread, so it was complemented by the hourly isochrones from the MTG FCI. In this case, ROS was calculated as the maximum distance in the wind direction between two consecutive hourly isochrones, as in previous publications (Duane et al., 2024). Statistical differences in ROS (in km h^-1^) between day and night were computed using analysis of variance.

To understand whether wildfires started burning quickly beyond capacity, we modeled the initial rate of spread, burned area over the first 8 hours, and the Initial Attack Assessment (IAA) using Wildfire Analyst (Cardil et al., 2025, 2023a). Fuels were derived from the Technosylva’s fuel map for the Iberian Peninsula, ignition points were provided by the local authorities, and weather conditions were obtained from Technosylva’s WRF model at 2 km pixel resolution. We used Wildfire Analyst for modeling because it has been thoroughly validated in prior studies (Cardil et al., 2025, 2023a). We compared the predicted ROS with measured ROS from VIIRS (r = 0.6) and sought expert opinion from the fire behavior analysts for further validation.

### Anomalies in fire weather and fuel moisture

For our analyses on environmental drivers, we examined patterns in live fuel moisture content (LFMC), weather (wind, temperature, and VPD), and the Fire Weather Index (FWI). Data on live fuel moisture content were obtained from previously developed products tailored specifically to the region of interest, with a 500m spatial resolution (Cunill Camprubí et al., 2022). That is, LFMC daily maps were computed using the LFMC-RF algorithm, which use MODIS MCD43A4 Nadir Bidirectional Reflectance Distribution Function (BRDF)-Adjusted Reflectance (NBAR) Collection 6.1 daily product (Schaaf & Wang, 2021) and MODIS MOD11A2/MYD11A2 Land Surface Temperature (LST) Collection 6.1 7-day composite products (Wan, 2014). Weather data, including 10m wind speed, 2m air temperature, 2m dew point temperature, and surface pressure, were obtained from ERA5-Land hourly data at 0.1° (Muñoz-Sabater et al., 2021). We report weather data at 15:00h on the day when the fires started.

Fire Weather Index data were downloaded from the Copernicus Emergency Management Service (https://ewds.climate.copernicus.eu/, last accessed on 12 May 2026), which provides daily data at a spatial resolution of 0.25 × 0.25° (Vitolo et al., 2020).

The analyses of anomalies in seasonal patterns of fuel moisture and weather were based on thresholds calculated at the daily scale using all data within an 11-day window centered on the target day. The thresholds are percentiles, which are appropriate for describing climate variability without making assumptions about the distributions of the variables used. That is, extreme values of 2025 represent the number or percentage of days below the 5th percentile, including those below the 0th percentile (when the absolute minimum was that recorded in 2025) within 2001-2024 for LFMC and DFMC. For weather variables (temperature, VPD, FWI, etc) extreme values were those at or above the 95^th^ percentile, including those above the 100th percentile (that is, when the absolute maximum in the time series occurred in 2025). Using an 11-day window allows for more robust threshold estimates due to a larger number of records, reducing the influence of skewed distributions caused by low data on a particular day, and facilitates the identification of intra-annual patterns (smoothing day-to-day variability) (Hobday et al., 2016; McDonald et al., 2018). We used the period 2001–2024 as the historical reference period for calculating weather statistics to match the fire database.

### Wildfire impacts on human health and ecosystems

To quantify wildfire impacts on human health, we quantified PM_2.5_ concentrations (PM_2.5_ derived solely from wildfires) from Zhang et al. (2025), and downloaded from https://tapdata.org.cn. Spatial population density was obtained from the WorldPop database (https://hub.worldpop.org/) using 2024 data 2024 at 1 km resolution from the R2025A v1 datasets of Portugal and Spain. WorldPop estimates are often normalized by official records (Láng-Ritter et al., 2025) from the United Nations (United Nations, Department of Economic and Social Affairs, Population Division, 2022) or, in the case of Europe, EUROSTAT (https://ec.europa.eu/eurostat/web/main/home). Following current approaches (Xu et al., 2023), grid-specific population counts within a country were normalized from EUROSTAT to ensure consistency with EUROSTAT census.

We estimated the mortality risk associated with wildfire PM_2.5_ from previously established approaches (Zhang et al., 2025). Specifically, daily deaths at the pixel level were calculated as:

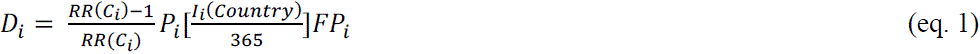

where *D*_i_ represents the estimated fatalities at pixel i; *RR*(*C*_i_) is the relative risk at PM_2.5_ exposure level *C* at the same grid *i*; *P*_i_ is the EUROSTAT-corrected population value; *I*_i_ is the baseline all-cause death rate for each specific Country; and *FP*_i_ is a conversion factor to translate the all-source PM_2.5_ mortality estimates to wildfire-specific mortality estimates following a direct proportion approach:

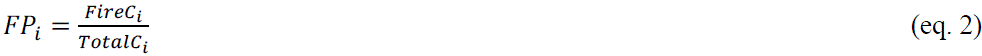

We derived RR from a recent meta-analysis (Alari et al., 2024) that estimated an increase of 0.08 in all-cause premature mortality per each 10 µg m^-3^ increase in PM_2.5_ exposure:

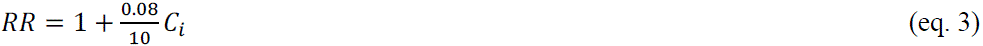

The baseline annual all-cause death rate (*I*) for Portugal and Spain was obtained from the Global Burden of Disease (GBD) (https://ghdx.healthdata.org/gbd-2023). The collected values were divided by 365 for a constant daily rate and converted to a raster map for calculations. After estimating daily premature deaths attributable to wildfires at a 10 km pixel level for the period of June-August 2025, we aggregated data at NUTS2 and NUTS3 boundary layers.

To characterize ecological wildfire impacts, we quantified burned area in protected areas. The protected areas layer came from the World Database on Protected Areas (WDPA), October 2025(UNEP-WCMC and IUCN, 2023), and was processed as in previously published studies (Resco De Dios et al., 2025). Land cover data were derived from country-specific land-use maps for Spain and Portugal: the Spanish Forest Map at 1:25,000 (MFE25, https://www.miteco.gob.es/es/cartografia-y-sig/ide/descargas/biodiversidad/mfe.html, last accessed on 12 May 2026) and the Carta de Ocupação do Solo Conjuntural – 2025 Pré-Verão for Portugal (https://snig.dgterritorio.gov.pt/, last accessed on 12 May 2026).

## Results and Discussion

### Increasing burned area and wildfire intensity

One of the key defining features of anthropogenic fire regimes over the last decades has been a decline in burned area (Andela et al., 2017). Southwestern Europe was no exception, and previous studies documented a decline in burned area since the 1980’s (Turco et al., 2016). This trend no longer holds after including the extreme wildfire seasons of 2022 and 2025. That is, the previously reported negative trend in burned area (Turco et al., 2016) becomes non-significant when considering the whole 1980-2025 trend (*p* = 0.16, simple regression, Fig. 1a). Instead, a segmented regression indicates a significant decline in burned area until 2022 (*p* = 0.03), and a positive (but non-significant) trend thereafter (Fig. 1a). This result is consistent with the “fire paradox” hypothesis, whereby initial declines in burned area resulting from fire suppression are not sustained over the long term due to large-scale fuel accumulations, which instead lead to more extreme fires and to more burned area (Minnich, 1983).

In addition to the burned area, changes in fire behavior could result from higher fire intensity. Geostationary data on Fire Radiative Energy (FRE) are available over a shorter period from SEVIRI (2003-2025) (Roberts et al., 2015). While no clear trend has emerged, 2025 showed the highest FRE recorded to date (26,339 TJ; Fig. 1b). To put this number in context, the total FRE emitted in 2025 by the Iberian Peninsula alone was similar to the annual energy production of a 1,000 MW nuclear reactor (∼30,000 TJ annually).

Most importantly, a significant increase in the mean Fire Radiative Power (FRP) has emerged in recent decades (Fig. 1c). Distinguishing between mean FRP and total FRE is essential, as the latter reflects both fire intensity and the number of hotspots (Schütze and Resco De Dios, 2025). Mean FRP, while controlling for hotspot detections, can serve as a quantitative proxy for fire intensity (Kaufman et al., 1996). Our analysis shows that, compared to the early 2000s, mild fire seasons now exhibit 40% higher fire intensities, and extreme seasons 30% higher fire intensities (Fig. 1c). That is, quantile regression of annual mean FRP revealed significant trends in both minimum (0.05) and maximum (0.95) quantiles. Minimum annual mean FRP rose from 35 to 49 MW per detection (40%) between 2004 and 2025, while maximum annual FRP increased from 63 to 82 MW per detection (30%). These findings indicate a major increase in wildfire intensity in the IP.

Recent studies on changes in fire behavior have reported increased nocturnal fire spread (Balch et al., 2022; Luo et al., 2024). We thus wanted to understand when such increases in annual mean FRP occurred. Nocturnal fire intensity rose marginally between 23.00-01.00h (*p* < 0.1) and significantly between 02:00-05:00h (*p* < 0.05, Fig. 1d), whereas changes during the rest of the day were not significant. Consequently, the observed annual increase in mean FRP (Fig. 1c) is primarily driven by intensified nighttime fire behavior, which has markedly strengthened over the 21^st^ century (Fig. 1d). These results highlight the growing role of nocturnal fire spread in shaping contemporary wildfire regimes.

### Increasing fire size and upward altitudinal expansion

The 2025 wildfire season was an extreme episode, largely confined to the western and northwestern parts of the Iberian Peninsula (Fig. 2a), with some important exceptions discussed below. According to the pyroregion classifications currently used in Spain (López Santalla and López García, 2019) and Portugal (Calheiros et al., 2020) (Fig. S7b), 43% of the burned area concentrated in the northwestern Spanish (NW-ES) pyroregion, 25% in southeastern Portugal (SE-PT), and an additional 17% occurred in NW-PT, with the remaining 15% distributed between Central Spain (C-ES, 13%) and Southeastern Spain (SE-ES, 2%). Across biomes (Dinerstein et al., 2017), 80% of the burned area occurred within the Mediterranean Forests, Woodlands and Scrub biome, and 20% in the Temperate Broadleaf and Mixed Forests biome. Burned area in NW-ES reached a record of 275,480 ha burned, 7.2 times above the 2001-2024 average (38,410 ha) and nearly double the previous maximum (147,729 ha, Fig. S1).

**Fig. 2.**
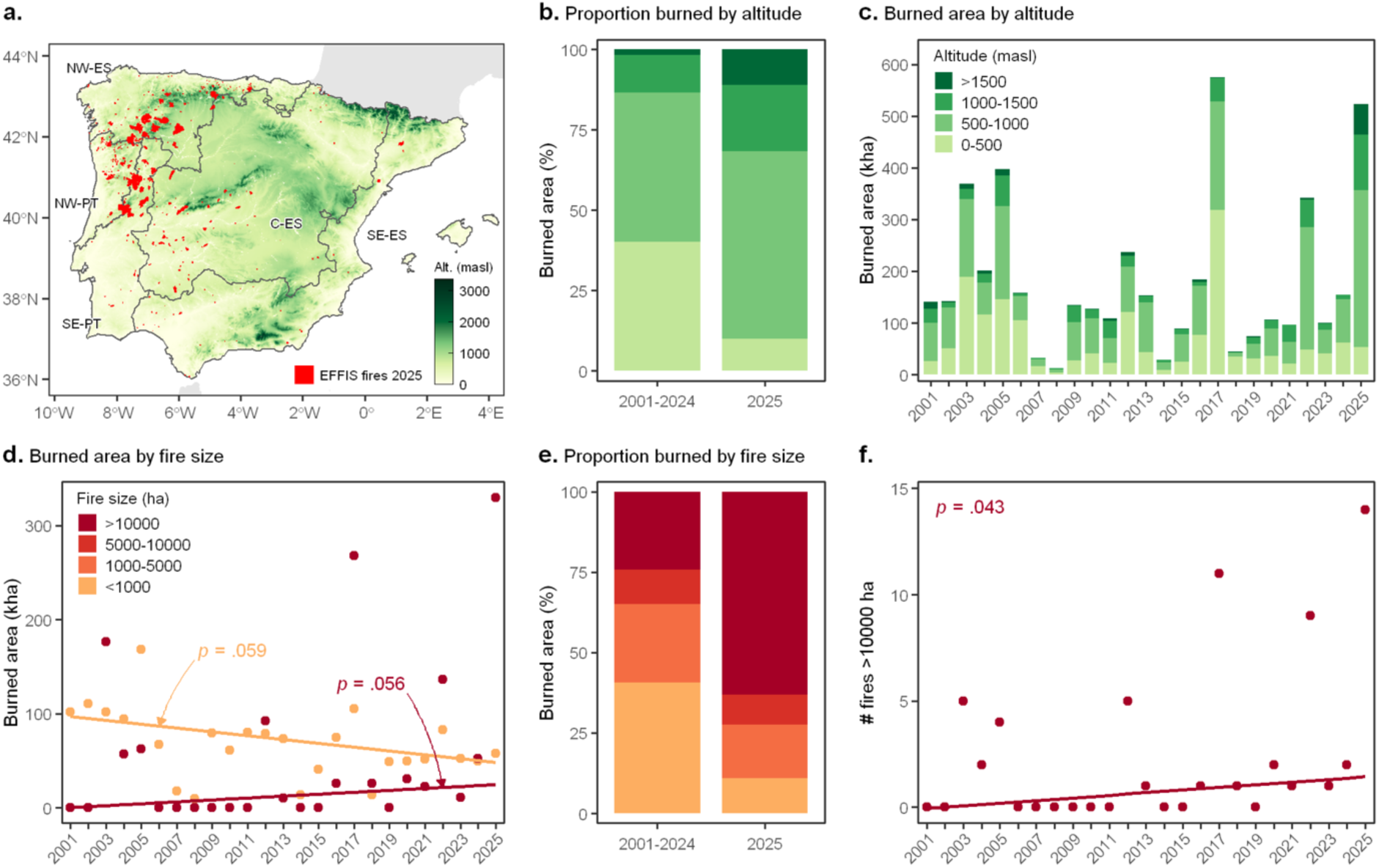
| Extremely large wildfires have been increasing in recent years and expanding into higher altitudes. (a) Spatial distribution of wildfires in 2025 across fire pyroregions; temporal changes in the (b) proportion and (c) total amount of burned area across different altitudes. The number of wildfires over 10,000 ha has increased marginally (d), and there have been significant increases in the proportion of burned area by large wildfires (e), and the total magnitude of wildfires over 10,000 ha (f).

A limited set of large fires drove the extreme burned area. According to EFFIS, there were 2,358 fires in the IP in 2025, of which only 102 large wildfires (> 500ha) were responsible for 94% of the total burned area and, within those, the 25 megafires (> 5,000ha) contributed 74% of the burned area (Fig. 2d-f, Table 1). This represents a shift in the relationship between fire-size distributions and total burned area relative to historical records. Total area burned by fires smaller than 1,000 ha has been decreasing since 2001 (from 91,751 ha to 42,865 ha on average, *p* = 0.059, Fig. 1d-e), whereas the contribution of fires larger than 10,000 ha has increased (from 9,195 ha to 95.355 ha on average, *p* = 0.056, Fig. 1d-e). The number of fires over 10,000 ha has also increased significantly (*p* = 0.043, Fig. 1f), reaching peak values in 2025, when they accounted for 63% of the total burned area, the highest proportion recorded to date (Fig. S2b). This pattern of “big fires getting bigger” is another indicator, albeit indirect, that fire intensity is increasing.

**Table 1.** | Main attributes for the 25 megafires (>5,000ha) that occurred during 2025 in the Iberian Peninsula. BA is the burned area, Land cover indicates the % cover for each vegetation type present in more than 20% of the burned area, VPD is the vapor pressure deficit, LFMC is the live moisture content, C-Haines is an index of atmospheric instability, and FWI is the fire weather index. All weather variables include maximum daily values. We additionally modeled the rate of spread (ROS) and burned area (BA) without fire suppression during the first 8 hours after ignition. IAA is the initial attack assessment.

| Date | Pyro-region | Fire | BA (ha) | Land cover | VPD (kPa) | LFMC (%) | C-HAINES | FWI | Modeled ROS (km h<sup>-1</sup>) | Modeled BA (ha in 8h) | IAA | Ignition source |
| --- | --- | --- | --- | --- | --- | --- | --- | --- | --- | --- | --- | --- |
| 1/7/2025 | SE-ES | Guissona | 5239 | Crops (76%)<br>Broadleaves (23%) | 5 | 73 | 11 | 48 | 2.94 | 6636 | 5 | Negligence /Accident |
| 26/7/2025 | NW-PT | Entre Ambos-os-Rios, Ermida e Germil | 7214 | Shrublands (71%) | 3 | 75 | 8 | 24 | 0.551 | 451 | 5 | Arson |
| 28/7/2025 | NW - PT | Real | 6546 | Eucalypts (69%) | 3 | 69 | 9 | 40 | 0.389 | 523 | 5 | Arson |
| 3/8/2025 | NW - PT | Pena, Quintã e Vila Cova | 6078 | Shrublands (55%)<br>Grasslands (22%) | 4 | 66 | 10 | 41 | 0.265 | 172 | 4 | Arson |
| 8/8/2025 | NW-ES | Manzaneda | 24418 | Shrublands (76%) | 3 | 73 | 10 | 45 | 0.446 | 330 | 5 | Arson |
| 8/8/2025 | NW-ES | Murias de Paredes | 7058 | Shrublands (71%)<br>Broadleaves (25%) | 1 | 73 | 10 | 45 | 0.351 | 388 | 5 | Lightning |
| 9/8/2025 | NW-ES | Benuza | 26199 | Shrublands (69%)<br>Broadleaves (21%) | 4 | 70 | 12 | 43 | 0.161 | 55 | 5 | Lightning |
| 10/8/2025 | SE-PT | Trancoso (São Pedro e Santa Maria) e Souto Maior | 62025 | Shrublands (37%)<br>Grasslands (33%) | 5 | 58 | 12 | 52 | 0.735 | 1562 | 3 | Negligence |
| 10/8/2025 | SE-PT | Sobral de São Miguel | 37578 | Shrublands (44%)<br>Conifers (25%) | 5 | 62 | 10 | 58 | 0.59 | 1387 | 5 | Lightning |
| 10/8/2025 | NW-ES | Uña de Quintana | 40018 | Broadleaves (33%)<br>Croplands (31%) | 5 | 60 | 12 | 55 | 0.975 | 3141 | 3 | Arson |
| 10/8/2025 | NW-PT | Ferreira de Aves | 13886 | Shrublands (46%)<br>Grasslands (23%) | 5 | 58 | 11 | 54 | 0.39 | 918 | 5 | Lightning |
| 10/8/2025 | NW-ES | Peranzanes | 7807 | Shrublands (79%) | 4 | 79 | 12 | 38 | 0.201 | 90 | 5 | Lightning |
| 11/8/2025 | NW-PT | Piódão | 30421 | Shrublands (60%) | 4 | 67 | 12 | 49 | 0.732 | 463 | 5 | Lightning |
| 12/8/2025 | NW-ES | Mezquita, A | 12008 | Shrublands (62%)<br>Conifers (21%) | 4 | 65 | 11 | 63 | 0.343 | 247 | 4 | Unknown |
| 13/8/2025 | NW-ES | Oímbra | 24434 | Shrublands (48%)<br>Conifers (27%) | 3 | 66 | 11 | 38 | 0.356 | 333 | 5 | Negligence<br>/Accident |
| 13/8/2025 | C-ES | Puertas | 10809 | Broadleaves (61%)<br>Grasslands (25%) | 4 | 52 | 11 | 46 | 0.500 | 453 | 3 | Lightning |
| 13/8/2025 | C-ES | Navaconcejo | 17298 | Shrublands (48%)<br>Broadleaves (37%) | 4 | 64 | 11 | 34 | 0.326 | 306 | 5 | Lightning |
| 14/8/2025 | NW-ES | Rúa, A | 37125 | Shrublands (57%)<br>Conifers (22%) | 4 | 77 | 11 | 38 | 0.189 | 130 | 3 | Unknown |
| 14/8/2025 | NW-ES | Burón | 19079 | Shrublands (72%)<br>Broadleaves (22%) | 2 | 68 | 8 | 20 | 0.179 | 206 | 3 | Lightning |
| 14/8/2025 | C-ES | Llerena | 5889 | Broadleaves (66%) | 5 | 51 | 12 | 63 | 0.75 | 1709 | 5 | Negligence /Accident |
| 15/8/2025 | NW-ES | Veiga, A | 24908 | Shrublands (78%) | 3 | 67 | 12 | 45 | 0.237 | 75 | 3 | Lightning |
| 15/8/2025 | SE-PT | Quintas de São Bartolomeu | 23567 | Grasslands (39%)<br>Shrublands (32%) | 5 | 60 | 11 | 63 | 0.858 | 1007 | 5 | Arson |
| 15/8/2025 | SE-PT | Alverca da Beira/Bouça Cova | 9512 | Shrublands (30%)<br>Grasslands (27%)<br>Croplands (25%) | 5 | 64 | 11 | 62 | 1.059 | 2500 | 5 | Arson |
| 15/8/2025 | SE-PT | Carviçais | 11916 | Shrublands (45%)<br>Grasslands (28%) | 5 | 64 | 12 | 58 | 0.532 | 881 | 3 | Negligence /Accident |
| 16/8/2025 | C-ES | Guardo | 6851 | Conifers (60%) | 4 | 70 | 13 | 50 | 0.75 | 1547 | 3 | Arson |

Nearly all the megafires (24 out of 25) started between 26 July and 16 August, and 84% (21 out of 25) in the first half of August (8-16 August, Table 1). These included the largest wildfires ever recorded in the Iberian Peninsula, like the Trancoso fire (PT) with 62,025 ha. The two largest wildfires on record in Spain also occurred in 2025: Uña de Quintana, with 40,018 ha, followed by Larouco, with 37,124 ha. It is worth noting that Galicia, one of the NUTS2 regions most affected by fire (Fig. S7) in NW-ES, had never experienced fires over 10,000 ha before 2022. However, five such fires occurred in 2025, ranging between 12,007 and 37,124 ha. Extreme fire behavior and instability led to 54 pyrocumulonimbus events across the IP, surpassing the previous record of 17 in 2022 by a factor of 3.1.

Another remarkable feature of the 2025 fire season was the extent of burned area at high altitude (Figs. 2a-c, S2a). Only 11% of the burned area occurred below 500m, the lowest proportion recorded during the observational period. Total burned area at 500-1000m, 1,000-1,500m, and above 1,500m was 1.5-, 2.5-, and 8.5-fold higher, respectively, than the peak values recorded during 2001-2024 (Fig. S2a). These values are consistent with previous projections (Alizadeh et al., 2021; Resco de Dios et al., 2021b) of an upward displacement of the fire belt under global warming, potentially reinforced by increased fuel loads from land abandonment.

### Conditions leading to the wildfires

The spring of 2025 was among the wettest since records began in 2001. Live fuel moisture content (LFMC) values in May were the highest recorded across large tracts of the IP (Fig. 3a-c, S3). Wildfires are associated with LFMC below 100% in forests and below 70% in shrublands (Rodrigues et al., 2023a), and LFMC was well above these thresholds across the IP in May (Figs. 3a-c, S3). Despite this wet spring, LFMC was below average in July and August 2025 (Figs. 3d-f, S3-S4), and fuels fell below critical LFMC thresholds, becoming readily available to burn. That is, there was a quick and strong dry-down of live fuels, particularly after the second half of July, which acted as a flash drought (Figs. S3-S4). However, record-breaking levels of fuel dryness were largely absent.

**Fig. 3.**
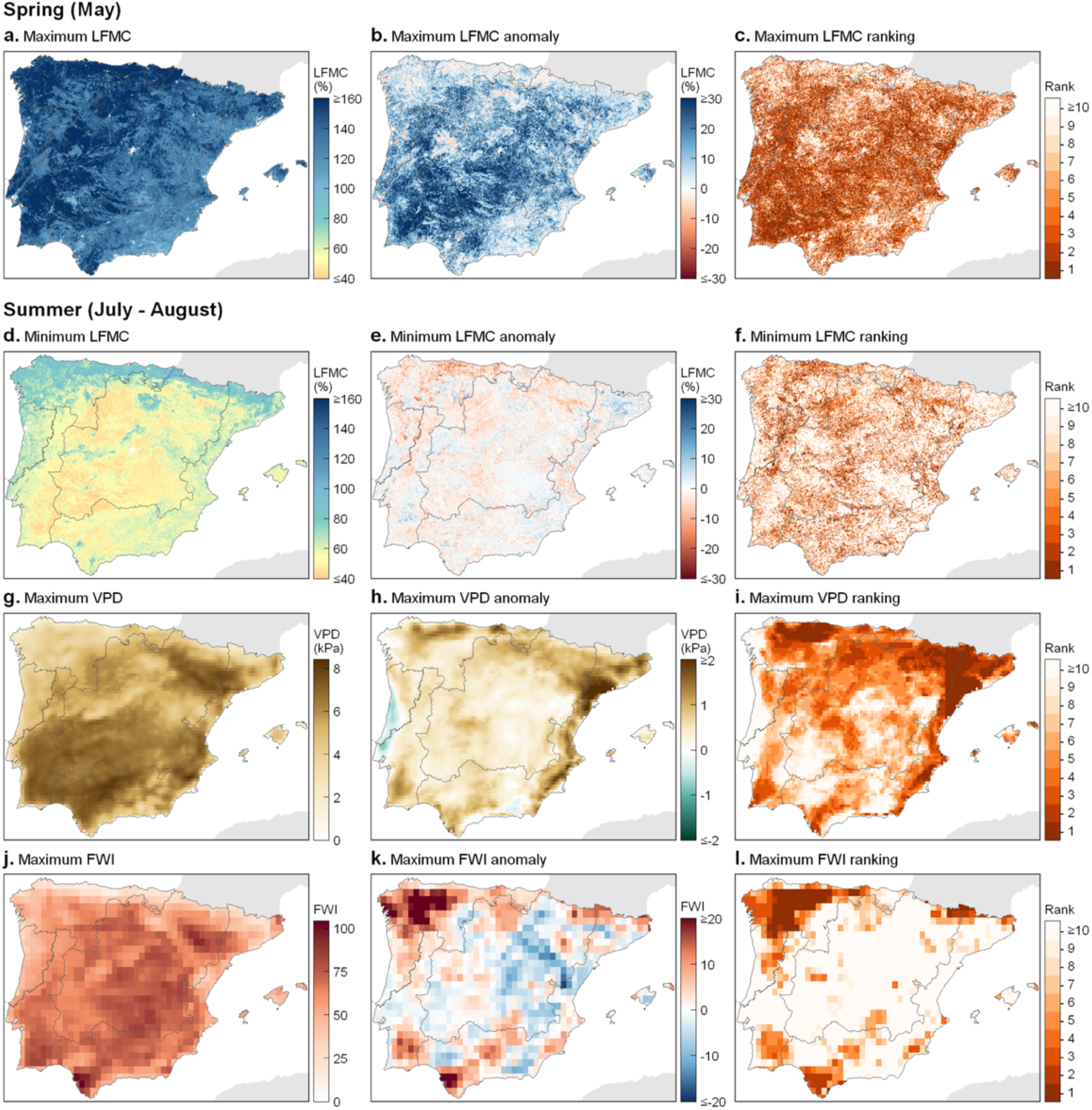
| Live fuel moisture content and fire weather during the 2025 wildfire season. Values of spring (May) live fuel moisture content (LFMC, a-c) and also of summer (July-August) LFMC (d-f), vapor pressure deficit (VPD, g-i), and fire weather index (FWI, j-l).

This contrasts with the 2022 fire season, the previous extreme fire year in southwestern Europe. In 2022, LFMC reached unprecedentedly low levels for 5–40% of the fire season, particularly in NW-PT, SW-PT, and C-ES (Rodrigues et al., 2023a). In 2025, however, extremely low LFMC values (below the 5th percentile) occurred only briefly (<5% of the summer) and were restricted to two pyroregions (SE-PT and NW-ES, Fig. S3). In other words, fuels were sufficiently dry to burn, but not exceptionally so (Figs. 3d-f, S3–S4).

In contrast, atmospheric drought intensified markedly. Vapor pressure deficit (VPD) increased sharply between June and early August, contributing to the rapid drying of both live and dead fuels (Fig. 2g–i, S3–S4). The Spanish Meteorological Agency reported that the summer of 2025 was the hottest on record, and the early August heatwave was the most intense in the observational series (AEMET, 2025). Beginning on 3 August, a 16-day heatwave exhibited a mean anomaly of 4.6 °C, exceeding the previous record set in 2022 (4.5 °C). Attribution analyses indicate that climate change made such an event approximately 40 times more likely, shifting its expected frequency from once every 500 years to approximately once every 13 years under current warming levels (Keeping et al., 2025). During the 2025 fire season, VPD exceeded the 95th percentile for approximately 20% of the July–August period, and record-breaking values were observed for a substantial fraction of the season (Fig. 2g–i, S3–S4).

**Fig. 4.**
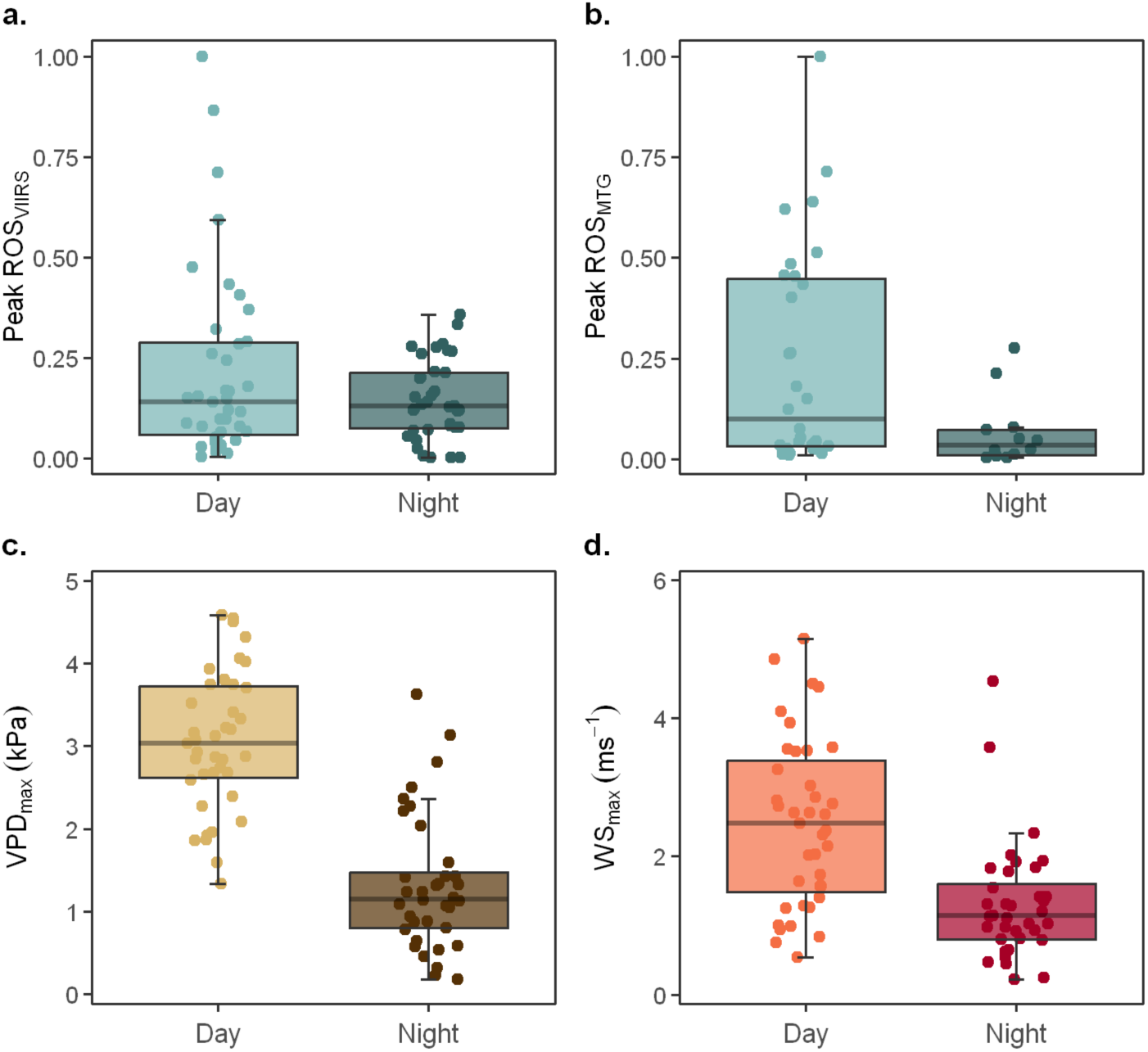
| Timing of maximum rate of spread (ROS). 58% of the wildfires occurring in the first fortnight of August showed peak ROS during daytime, while ROS peaked at night in the remaining 42% according to VIIRS (a). This proportion changed to 71-29% using fire isochrones derived from Meteosat Third Generation (b, MTG). However, maximum vapor pressure deficit (VPD) and wind speed (WS) were always higher during the day than at night, indicating that surface meteorological conditions alone cannot explain this pattern. ROS is presented in relative units (after linear rescaling) to accommodate differences in both methods to estimate ROS.

We also observed a marked decoupling between fuel moisture and fire weather conditions. While LFMC was not exceptionally low, fire weather indices were extreme across large parts of NW-ES (Fig. 3j–l, S3–S4). In particular, the Initial Spread Index (ISI) and the Build-up Index (BUI) exceeded the 95th percentile for more than 40% and 75% of the summer, respectively (Fig. S3–S4). The Fire Weather Index (FWI) also reached extreme values in other regions affected by megafires, including NW-PT, SE-PT, and C-ES, although over shorter durations and smaller spatial extents.

### Fuel status in the megafires

To better understand the 2025 fire season, we examined different environmental factors associated with the 25 megafires. As previously noted, forests and shrublands in southwestern Europe start to burn as LFMC drops below 100% and 70%, respectively (Rodrigues et al., 2023a). Across the 25 megafires, the mean LFMC was 66% (range 50-78%). These values are characteristic of shrubby fuels and do not represent extreme dryness, as shrublands can exhibit LFMC values below 50% (Pellizzaro et al., 2007) (Table 1).

Large wildfires have been associated with DFMC values below 10% (Nolan et al., 2016). Vapor pressure deficit exceeded 3 kPa in most cases, which, according to the model of Rodrigues et al. (2024), corresponds to a dead fuel moisture content below 7%. Similarly, the Fire Weather Index (FWI) was above the critical value of 39 (Alexander and de Groot, 1988), associated with fire intensities above 10,000 kW m^-1^ (the limit for suppression capacity (Tedim et al., 2018)), although higher values were also recorded in 2022 (Rodrigues et al., 2023a).

Vegetation burned during these megafires was predominantly shrubby (Fig. S5, Table 1), with 46% of the burned area corresponding to shrublands, followed by native broadleaves (17%) and conifer forests (10%). The category “native broadleaves” is not limited to forests; it also includes savanna-like *dehesas* and *montados*. Other land uses, such as crops or Eucalypt plantations, accounted for less than 10% of the burned area. Although not captured by current vegetation inventory protocols, substantial grass growth may have occurred after the wet spring, subsequently curing during the summer dry-down (Muñoz-Gómez et al., 2024). This could have increased the availability of fine dead fuels, with a potential yet unquantified role in increasing fire spread rates and spotting activity.

### Nocturnal fire acceleration played a key role

A defining feature of this wildfire season was the prominent role of nighttime propagation (Fig. 4a-b). We calculated the rate of spread (ROS) during the peak fire season (first half of August) using two independent approaches: burned area from VIIRS (Cardil et al., 2023b) and Fire Radiative Power from the geostationary Meteosat Third Generation (Xu et al., 2026) as explained in the methods section. Both approaches indicate a substantial contribution of nocturnal fire spread, with peak ROS occurring overnight in **29–42% of wildfires** (depending on the method; Fig. 4a-b). In other words, fire spread rates were higher at night than during the day in up to 42% of cases. When peak ROS occurred at night, values were not significantly different from daytime peaks, based on VIIRS estimates (*p* > 0.1).

Previous studies have documented increasing nocturnal fire spread in recent years (Balch et al., 2022; Luo et al., 2024), consistent with the disproportionate warming of nighttime temperatures under climate change. VPD is a major driver in fire activity(Clarke et al., 2022), and climate change-driven increases in nocturnal VPD are thus expected to increase fire spread overnight.

However, here we show that wildfires burned faster at night than during the day in at least 29% of cases, and that nighttime peak ROS values were not lower than daytime values when using VIIRS (Fig. 4). This is more difficult to explain because, even if nocturnal VPD has increased, daytime VPD will still be higher than nocturnal VPD. In fact, we observed that both VPD and wind speed were significantly higher during the day than during the night (Figs. 4c-d). In other words, changes in near-surface conditions cannot fully account for the observed nocturnal maxima in ROS. Such behavior has been seldom observed (Castellnou et al., 2024; Luo et al., 2026). In North America, a recent study documented that FRP peaked at night in 14% of active fire days, a frequency 50% lower than that reported here (Luo et al., 2026). A previous study (Castellnou et al., 2024) hypothesized that wildfires may burn faster at night due to pyroconvection, when fire-induced plumes penetrate into the free troposphere, deepening the atmospheric boundary layer and accelerating fire spread (Castellnou et al., 2024). Consistent with this mechanism, the Continuous Haines index (C-Haines), an indicator of atmospheric instability (Mills and McCaw, 2010) previously validated in this region (Resco de Dios et al., 2021a), exhibited elevated values across most megafires (Table 1). Future research efforts should concentrate on understanding the mechanisms underlying this behavior.

### Lightning was the main ignition source

From the 25 megafires, lightning was the ignition source in at least 10 (40%). Arson started 8 megafires (32%), and negligence or accidents were responsible for a further 5 ignitions (20%). At the time of writing, the cause of ignition for 3 megafires remained unknown (Table 1, Fig. S6).

These results are important because arson is often emphasized as a dominant ignition cause in megafires in public discourse, and it is often stated that 95% of all ignitions in SW Europe are anthropogenic (San-Miguel-Ayanz et al., 2012). While the latter statement is true, it is incomplete, as lightning is often the dominant ignition source in megafires. Considering only the 16 largest fires, those over 10,000ha, lightning ignited 50%, and this percentage rose to 70% in the 2022 fire season (Rodrigues et al., 2023a). Although lightning accounts for only a small fraction of total ignitions, it contributes disproportionately to the largest megafires. Human ignitions typically occur near infrastructure (Costafreda-Aumedes et al., 2018; Gonzalez-Olabarria et al., 2012) and are more accessible to suppression teams, whereas lightning ignitions occur more randomly, often in remote, mountainous areas that are more difficult to access (Rodrigues et al., 2023b). Lightning often occurs during dry storms, which can enhance fire spread because they are associated with higher wind speeds and atmospheric instability (Pérez-Invernón et al., 2023).

### Fire simultaneity was large, but not extreme

We tested whether an unprecedented level of fire simultaneity contributed to the 2025 season by examining the relationship between ignition numbers and burned area. Overall, we did not find support for the hypothesis that the 2025 fire season was driven primarily by an exceptional number of simultaneous ignitions (Fig. 5). Across all regions, there were 5,185 ignitions in 2025, a value below the regional median of 8,146 ignitions (in the 0.28 quantile; Fig. 5a) and well below historical maxima, as over 20,000 ignitions were recorded in 1998 (29,291 ignitions), 1995 (22,865 ignitions), and 2005 (20,253 ignitions). The number of ignitions was towards the higher end in some regions (Castilla y León was in the 0.72 quantile, Figs. 5e, S7), but not in others (Centro or Norte were in the 0.13 and 0.33 quantiles, respectively, Fig. 5b,c), relative to the historical record. In other words, the burned area in 2025 was substantially higher than would be expected from the historical relationship between ignition numbers and burned area.

**Fig. 5.**
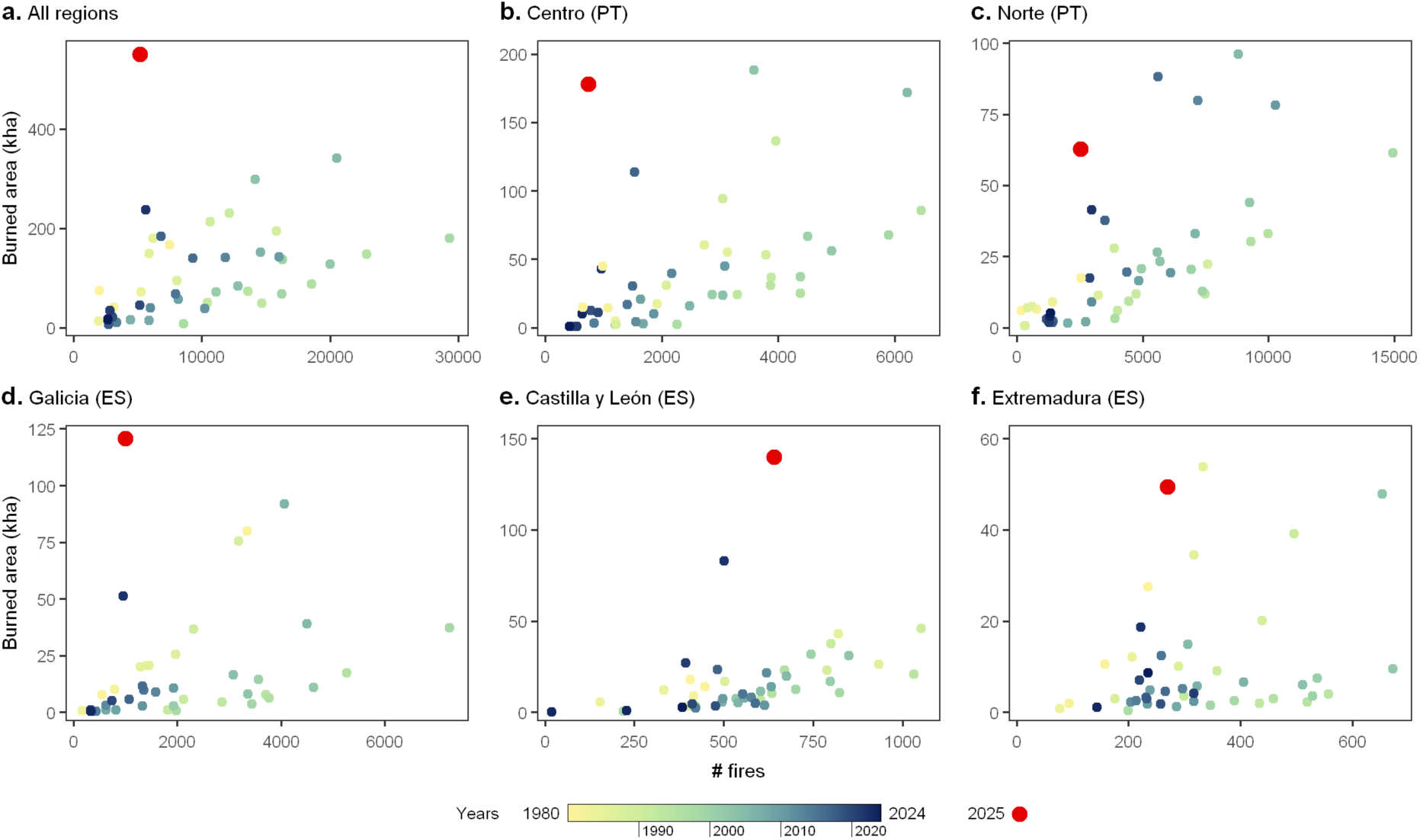
| Fire simultaneity alone cannot explain the large burned area. Relationship between the number of ignitions during the fire season (July-August) since 1980 in the 5 most affected NUTS2 regions (Fig. S7) from Spain (Castilla y León, Galicia, and Extremadura) and Portugal (Norte and Centro). Colors indicate year, with yellow representing early years in the dataset and blue representing recent years. The year 2025 is shown by the bigger red dot. The number of wildfires documented during this period was not exceptionally high relative to previous wildfire seasons.

### Firefighting was likely overwhelmed, allowing fires under low initial spread to become megafires

Fire simultaneity could still contribute to the 2025 season if global change has increased our vulnerability to concurrent events. This could occur under increasing fire intensities: initial fires require so many resources that subsequent ignitions are left unattended, allowing them to develop beyond capacity (Petrovic et al., 2012; Thompson et al., 2023). Although fire suppression capacity has increased in the Iberian Peninsula in recent decades (Ortega et al., 2026; Urbieta et al., 2019), increasing fuel accumulation under a warming climate could increase vulnerability to simultaneous fires.

We thus modeled initial attack assessment (IAA), ROS, and burned area over the first 8 hours (Cardil et al., 2025). IAA is a categorical index (1-5), where higher values indicate a declining probability of successful suppression (IAA=5 corresponds to a 90% decrease in initial attack success odds) (Cardil et al., 2025). Based on previous work (Cardil et al., 2025), we considered an initial attack likely ineffective under IAA ≥ 3, mean ROS > 300m h^-1,^ and an estimated burned area exceeding 500 ha within the first 8 hours. Only 9 fires met all three criteria (Table 1). In contrast, 6 megafires showed initial ROS below 300 m h^-1^ and, consequently, were not expected to exhibit extreme fire behavior during the initial 8 hours.

Overall, 24% of all megafires started with non-extreme behavior, indicating that suppression may have been effective with a strong initial attack (Table 1). Their eventual growth is consistent with suppression teams being overwhelmed, particularly given the strong spatial and temporal clustering of these wildfire events.

We hypothesize that the primary control on overwhelming suppression teams is the cumulative perimeter of simultaneous fires, rather than the number of ignitions *per se*. During the first half of August, cumulative fire perimeters reached 1,529 km in Castilla y León and 1,312 km in Galicia, indicating an extreme, spatially clustered suppression demand. Such an extent makes suppression saturation plausible, even in well-resourced systems. Operational constraints, including the prioritization of settlements and infrastructure, may further reduce suppression efficiency. Despite increasing investment in fire suppression (Urbieta et al., 2019), system effectiveness may be declining under global change, as rising fire intensity reduces the success of initial attacks, and increasingly extreme fire weather enhances fire spread and expands concurrent fire perimeters.

### Impacts

We calculated the expected premature deaths resulting from the PM_2.5_ emissions. Cumulative PM_2.5_ concentrations varied widely across Spain, reaching values over 400 mg m^-3^ in the areas closer to the wildfires (Fig. 6a, S8). Based on established exposure-response functions (Chen et al., 2021), these increases in PM_2.5_ are expected to have led to 2,065 chronic premature fatalities (Fig. 6b, S8) among individuals with pre-existing conditions, including 443 in Portugal and 1,622 in Spain. The region of Madrid suffered the largest expected mortality values, with 359 fatalities, followed by Catalonia (268 fatalities) and Andalusia (200 fatalities).

**Figure 6.**
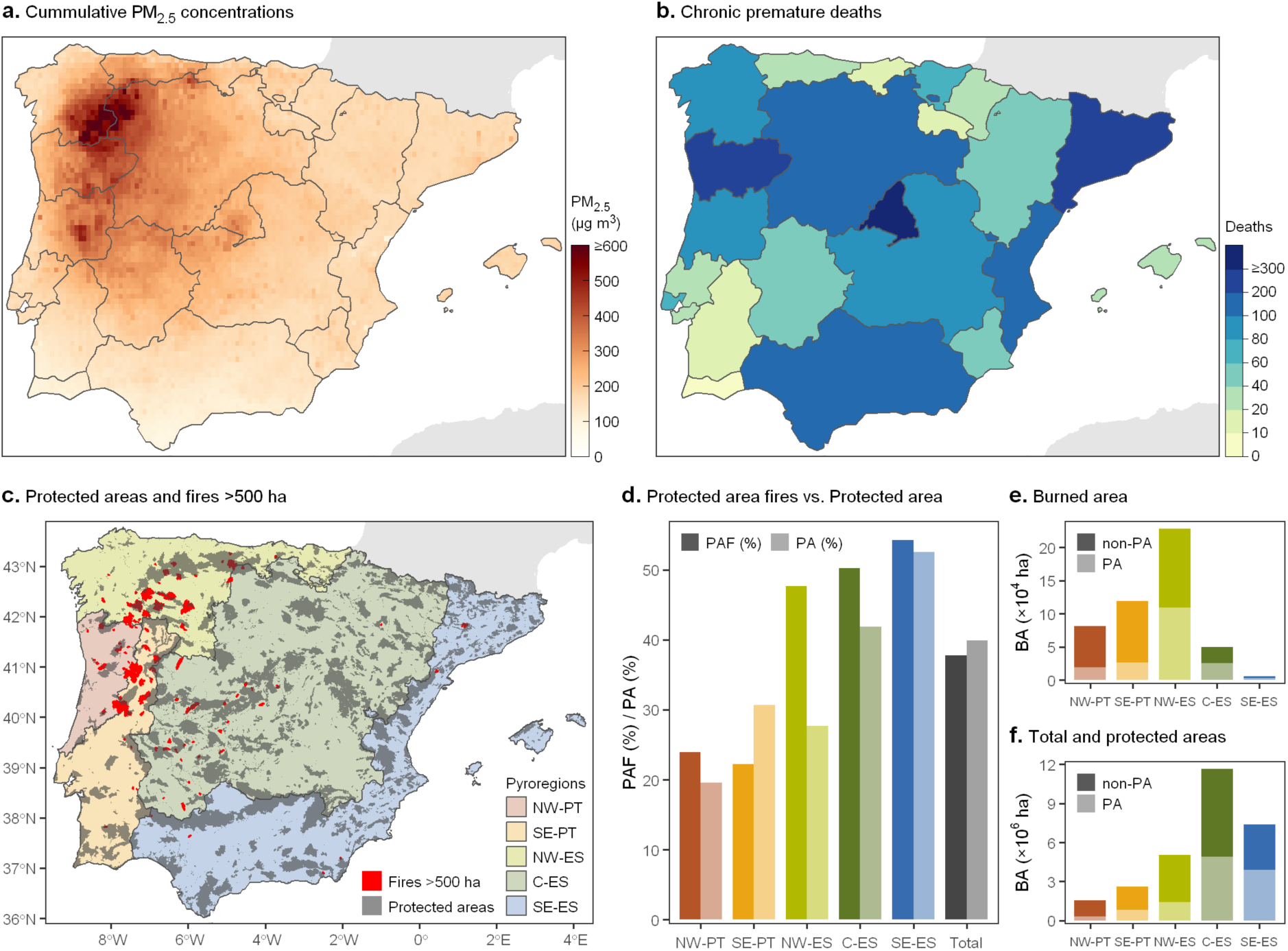
| Wildfires increased PM_2.5_ and associated mortality and also impacted protected areas significantly. We mapped PM_2.5_ emissions from wildfire (a) and calculated expected premature deaths (b) across NUTS2 regions in the Iberian Peninsula (Fig. S7). We also assessed whether impacts on protected areas were disproportionate across pyroregions (c, Fig. S7), by comparing the proportion (d) of the wildland area that is protected (PA, shown in bright colors), with the proportion of the total wildland area burned in large wildfires (> 500ha) occurring within protected areas (protected area fire or PAF, in shaded color). Additionally, total burned area in large fires (e) across protected and unprotected areas (PA and non-PA, respectively); and the distribution of the cover of protected (PA) and unprotected (non-PA) in wildlands (f).

In addition to these chronic deaths, official reports indicate 14 fatalities as a direct result of wildfires in the Iberian Peninsula, 8 in Spain (DG Protección Civil, 2026), and 6 in Portugal (SGIFR and AGIf, 2026). A further 65 people were injured in Spain, of whom 55% were fire-fighting personnel and 45% were civilians. Causes of injuries were varied, ranging from operational incidents (e.g. vehicle accidents or during mop-up operations), as well as smoke inhalation or severe burns. A total of 39,405 evacuees were recorded in Spain, most of them (28,368) concentrated in Castilla y León (DG Protección Civil, 2026). Sixteen wildfires affected infrastructure, including towns and industrial areas, and over 70 wildfires impacted transport networks (roads and rails). Firefighters prioritize population protection, which partly explains why some wildfires in remote areas were left unattended during peak activity periods. Official data on injuries, evacuations, and infrastructure impacts were not available from the Portuguese civil protection service at the time of writing.

Considering impacts on wildlands (that is, excluding crops, urban areas, and water bodies), we quantified that 183,116 ha of protected areas (PAs) were burned (Fig. 6c-f), which is equivalent to 38% of the total wildland area. Protected areas cover 40% of the total wildland area, so they were not disproportionately affected by fire. However, impacts varied geographically. PAs burned disproportionally in the NW-ES pyroregion, as PAs only occupy 27.7% of the wildland area, but concentrated 47.7% of the total area burned. In NW-PT and C-ES, the disproportionate impact was smaller, at 4% and 8%, respectively. In contrast, in SE-PT, protected areas burned less than expected, as they occupy 31% of the wildland area but account for only 22% of the burned area. These results are consistent with a brief report focusing on NW IP, which similarly identified a disproportionate impact on protected areas NW-PT and C-ES(Beltrán-Marcos et al., 2026).

Protected areas affected by fire include national parks that are home to emblematic species like capercaillie (*Tetrao urogallus* L.) or wolf (*Canis lupus* L.). Wildfire impacts on fauna, however, are species-specific, and they have been reported to be positive for apex predators such as wolves(Lino et al., 2019), and expected to be negative for some other species. From an ecological perspective, burned area alone cannot be directly equated with negative ecological impact, as outcomes depend on the spatial configuration of burned patches, fire severity mosaics, recurrence intervals, and species-specific fire sensitivity(Jones and Tingley, 2022). Unraveling the ecological consequences of the 2025 fire season will require longer-term monitoring and follow-up studies (Driscoll et al., 2024).

### Future outlook

The 2025 wildfire season likely provides a potential preview of future fire activity under unabated climate change and land abandonment. Longer time series will be needed to confirm whether we have entered a new fire regime, but we quantify major changes in key underlying components. Fire intensity has increased by 30-40% during the 21^st^ century, primarily due to nighttime fire behavior, and record-breaking levels of mean FRP and total FRE were achieved in 2025. Nocturnal wildfire intensity has increased to the point where 29-42% of all wildfires now spread faster at night than during the day, indicating an unexpected acceleration at night that substantially narrows the operational window for fire suppression.

The extreme fire season of 2025 in the Iberian Peninsula resulted from the interaction of multiple factors. Predisposing factors included low live fuel moisture content, high vapor pressure deficit, and extreme fire weather during the early August heatwave. The 2025 summer heatwaves behaved like flash droughts, quickly drying fuels beyond critical dryness thresholds, despite a preceding very wet spring. However, these factors alone cannot fully explain the observed fire behavior, particularly the nocturnal peaks in fire spread. Nocturnal wildfire acceleration cannot be explained solely by surface weather and fuels: VPD and wind speed were lower at night. Pyroconvection and fire coupling with the free troposphere may have contributed to driving the extreme fire behavior observed (Castellnou et al., 2024), a poorly understood process that should be at the forefront of our research efforts. Atmospheric instability, coupled with lightning activity starting on August 8^th^, contributed to a simultaneous fire event. After that point, our results are consistent with a saturation of firefighting systems, as very large cumulative fire perimeters became unmanageable, further exacerbating wildfire spread.

Most megafires ignited before the 8^th^ of August showed a low likelihood of successful initial attack. The megafires that started after that date, however, comprised a combination of wildfires that may have been controllable under an intense initial attack and others that quickly exceeded capacity. The rapid expansion of fire perimeters after the 8^th^ of August likely overwhelmed firefighting systems, allowing some initially manageable fires to develop into megafires.

Limitations in firefighting capacity may have contributed to the severity of the season due to an increase in fire intensity. These results indicate that bigger investments in fire prevention are needed, especially as the temporal window for fire suppression is narrowing.

Anthropogenic fire regimes during the 20^th^ century were characterized by declining burned area due to substantial investments in fire suppression (Archibald et al., 2013). In contrast, emerging patterns in the 21st century indicate increasing fire size and simultaneity globally. For example, 21% of the temperate broadleaf forest biome in southeastern Australia burned during the 2019–2020 season (Boer et al., 2020), and approximately 13 million hectares burned in Canada in 2023 (Jain et al., 2024). This evolving regime also includes an upward expansion of fire activity into previously less affected areas, as documented here.

Fire suppression may become less effective under global change, increasing the importance of preventive measures. As climate change intensifies, fire prevention will need to be implemented under increasingly larger areas, and delays in preventive action today are likely to increase future costs.

## CRediT authorship contribution statement

Víctor Resco de Dios: Conceptualization, Formal analysis, Funding acquisition, Investigation, Project administration, Writing – original draft. Àngel Cunill Camprubí: Formal analysis. Simon J Schütze: Formal analysis, Writing – review and editing. Fernando Castedo-Dorado: Conceptualization, Formal analysis, Writing – review and editing, Juan Picos: Conceptualization, Writing – review and editing, Joaquín Ramírez: Conceptualization, Formal analysis. Rut Domènech: Conceptualization, Writing – review and editing. Mercedes Bachsfischer: Formal analysis, Writing – review and editing. Marc Castellnou: Conceptualization, Writing – review and editing. Adrián Cardil: Conceptualization, Formal analysis, Writing – review and editing.

## Data availability

This manuscript uses publicly available datasets as referenced in the methods

## Acknowledgements

We acknowledge funding from the Spanish MICINN (PID2022-138158OB-I00).

## Supplementary Information for

**Fig. S1.**
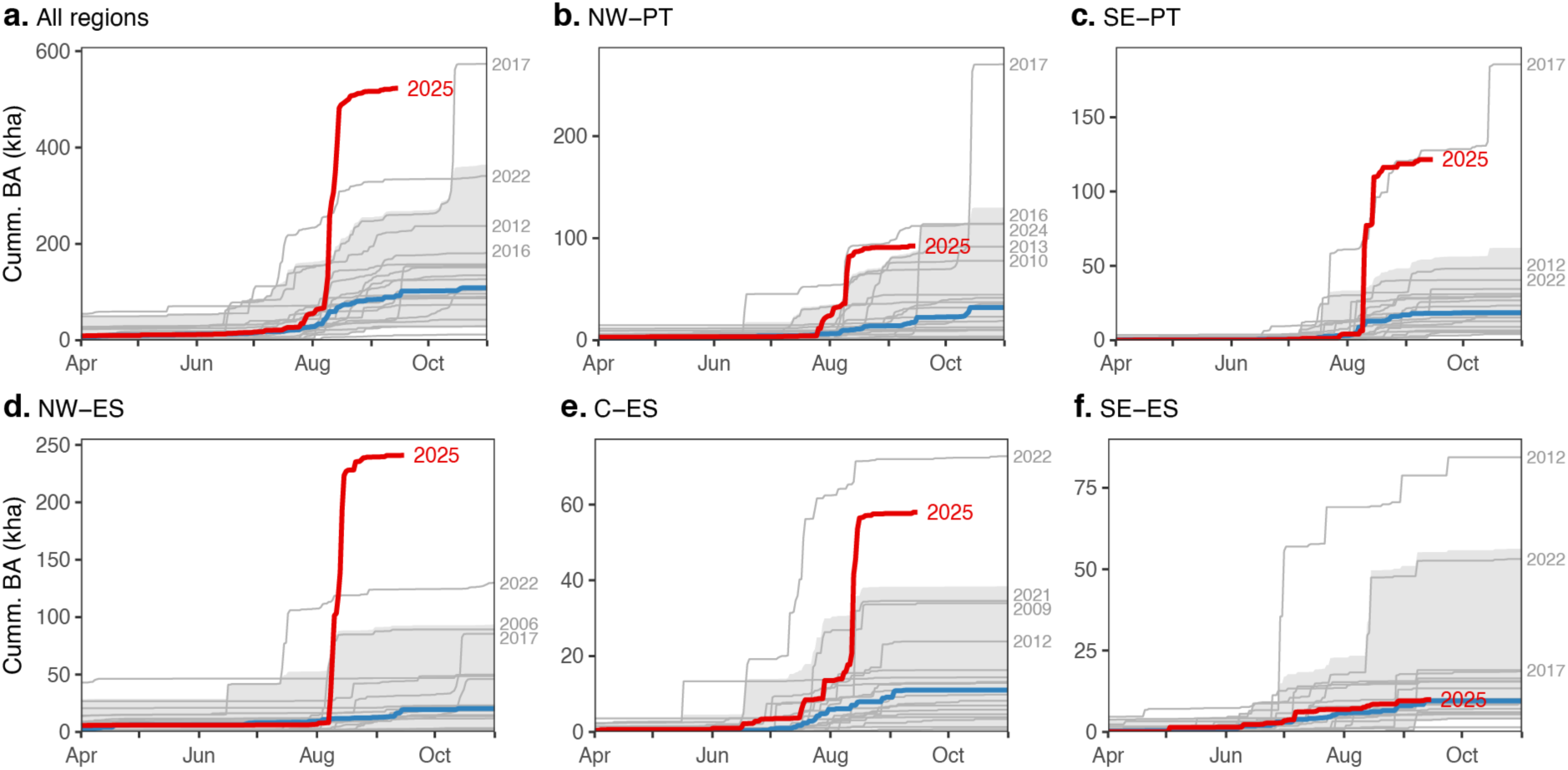
| Cumulative burned area in SW Europe. The shaded area indicates the 95th percentile in the historical records (2001−2024) and the blue and redlines the historical median and 2025, respectively.

**Fig. S2.**
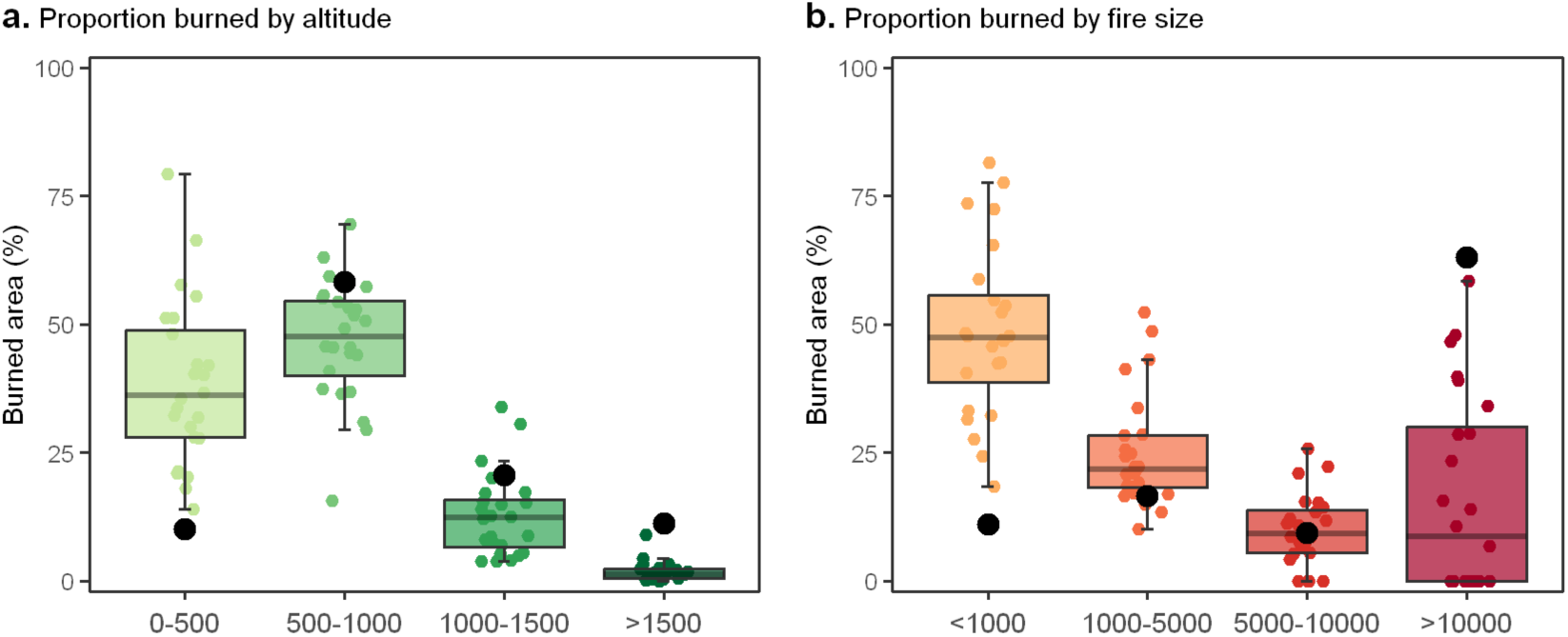
| The proportion of burned area across 2001-2025 as a function of altitude (a) and fire size (b).

**Fig. S3.**
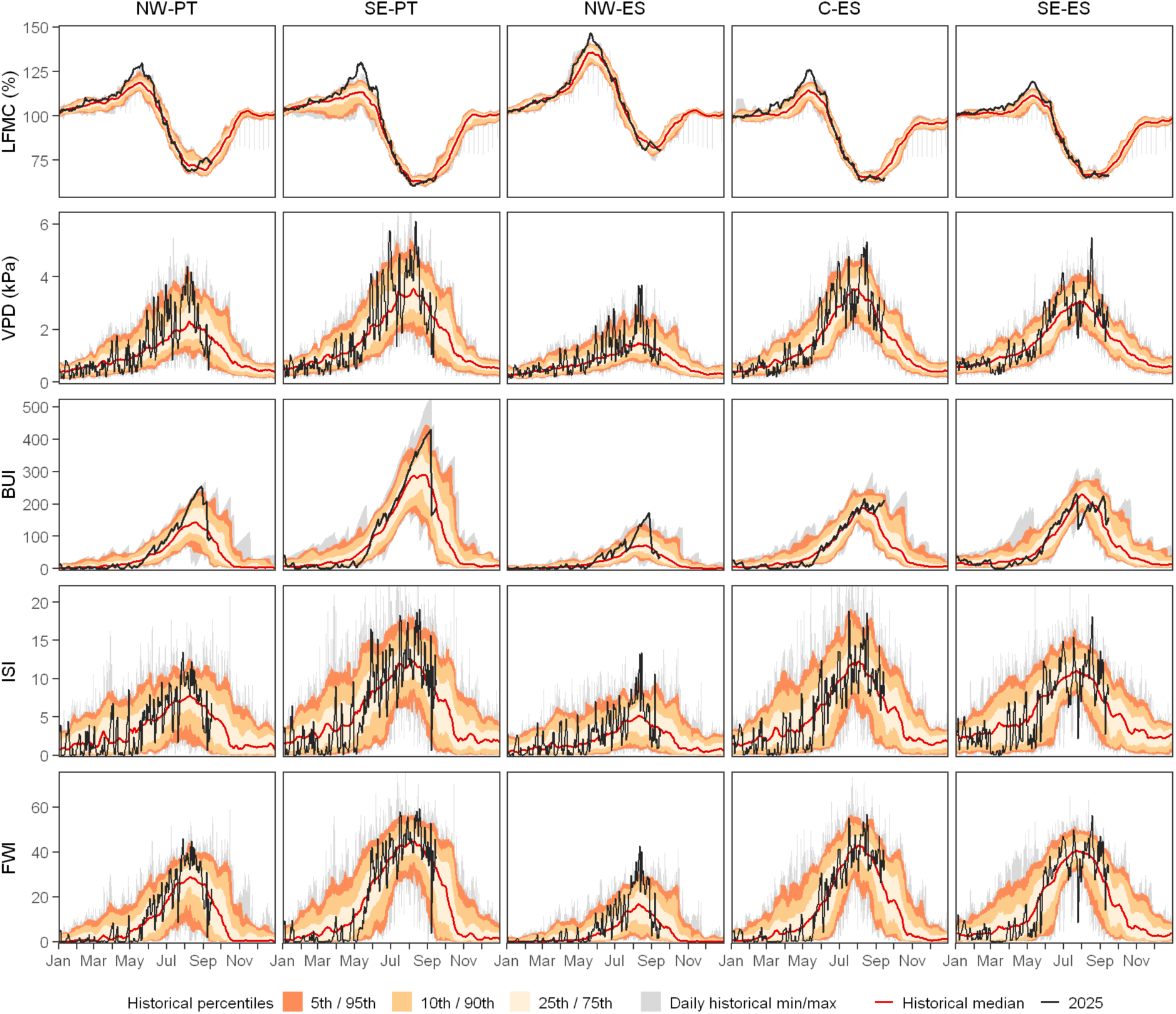
|Temporal evolution of live fuel moisture content (LFMC), vapor pressure deficit (VPD), Initial Spread Index (ISI), Build-up Index (BUI), and Fire Weather Index (FWI) across the different pyroregions in 2025 in relation to the period 2001-2024. The LFMC peak in May was the highest recorded during the study period, followed by very high VPD values, which acted like a flash drought, and with very high fire weather.

**Fig. S4.**
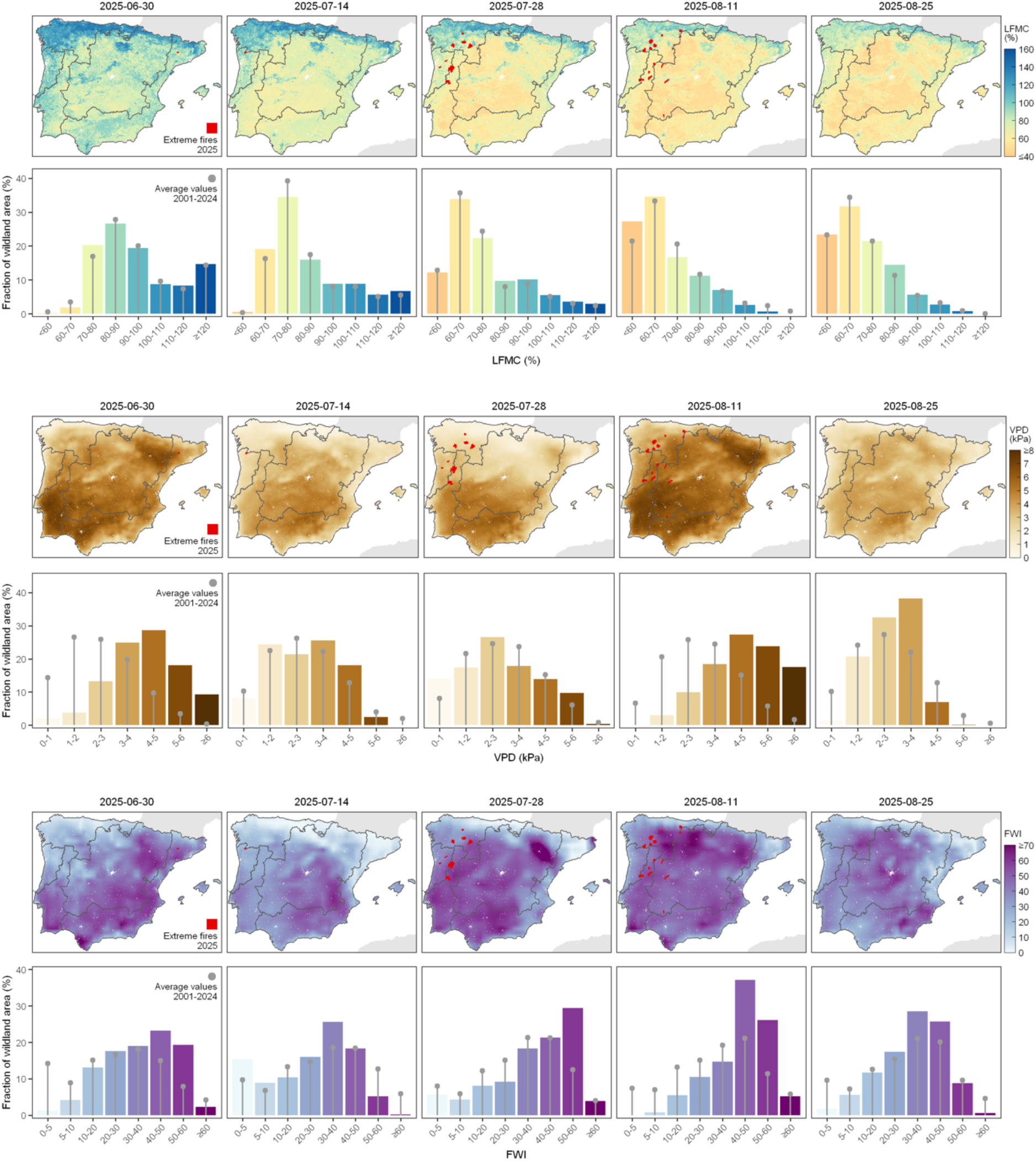
| Temporal evolution of fuel moisture and fire weather. The graphs depict the progression of changes in live fuel moisture content (LFMC), vapor pressure deficit (VPD) and fire weather index (FWI).

**Fig. S5.**
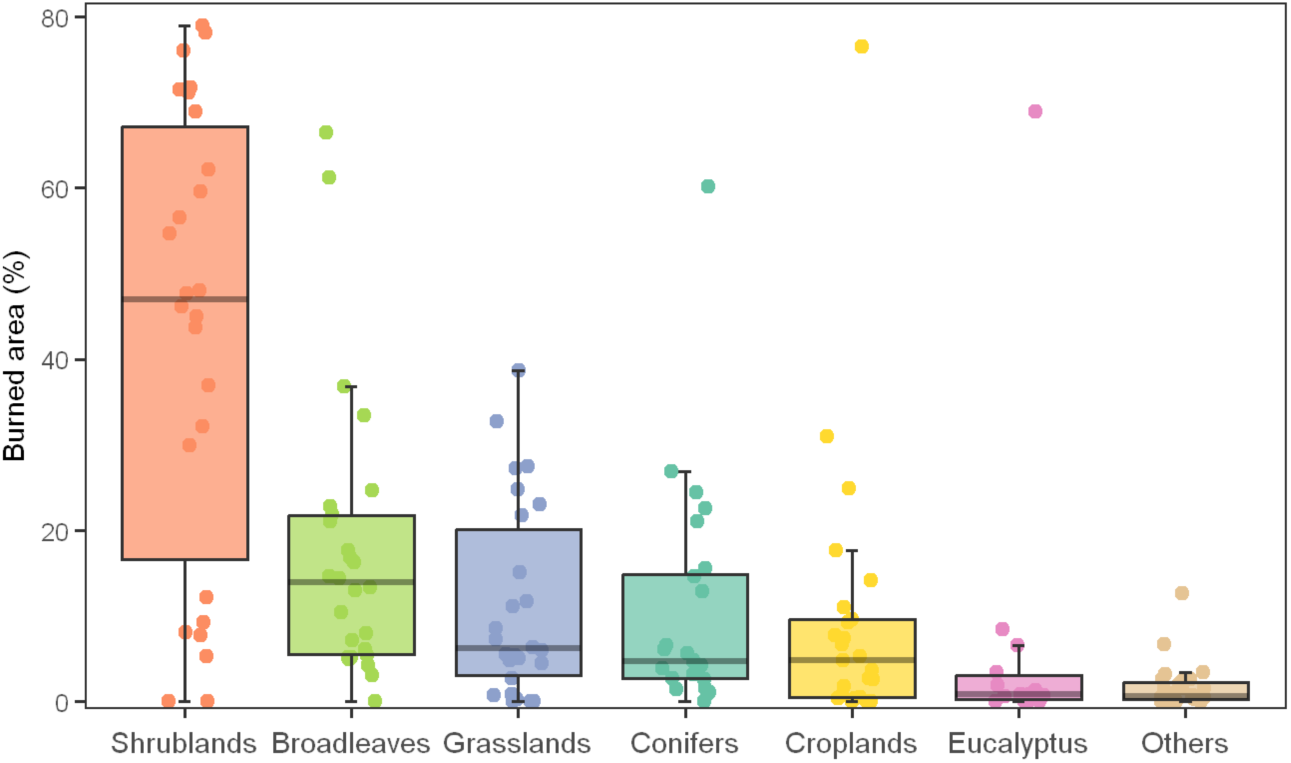
– Burned area across vegetation types in the 25 megafires. Red diamonds indicate average cover in the area.

**Fig. S6.**
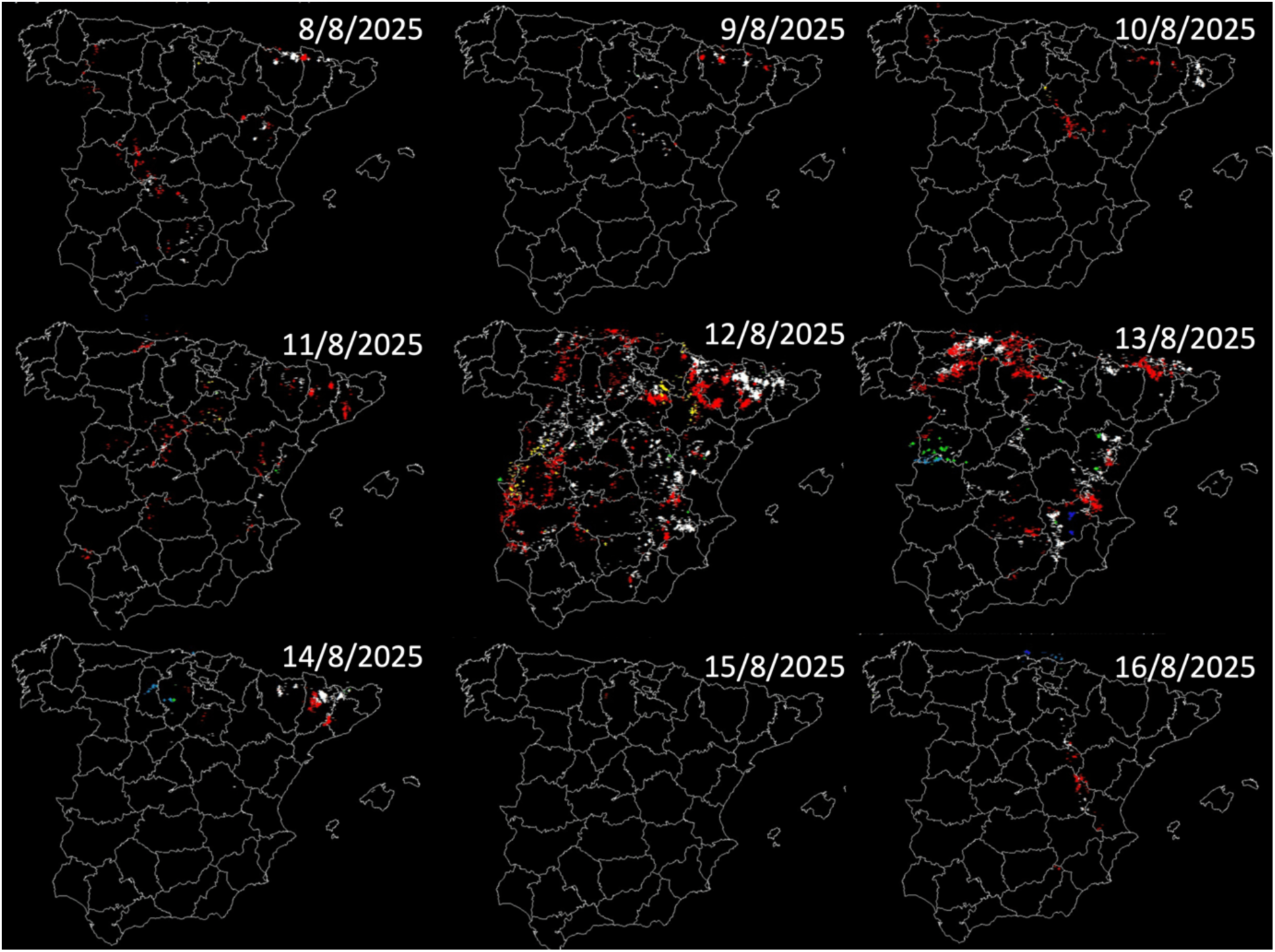
| Lightning activity between 8 and 16 August 2025 in continental Spain. White, blue, green, white, red, and yellow colors indicate 00:00:01h-04:00:00h, 04:00:01h-08:00:00h, 08:00:01h-12:00:00h, 12:00:01h-16:00:00h, 16:00:01h-20:00:00h, 20:00:01h-00:00:00h, respectively. Source: https://aemetblog.es (accessed on 12^th^ May 2026).

**Figure S7.**
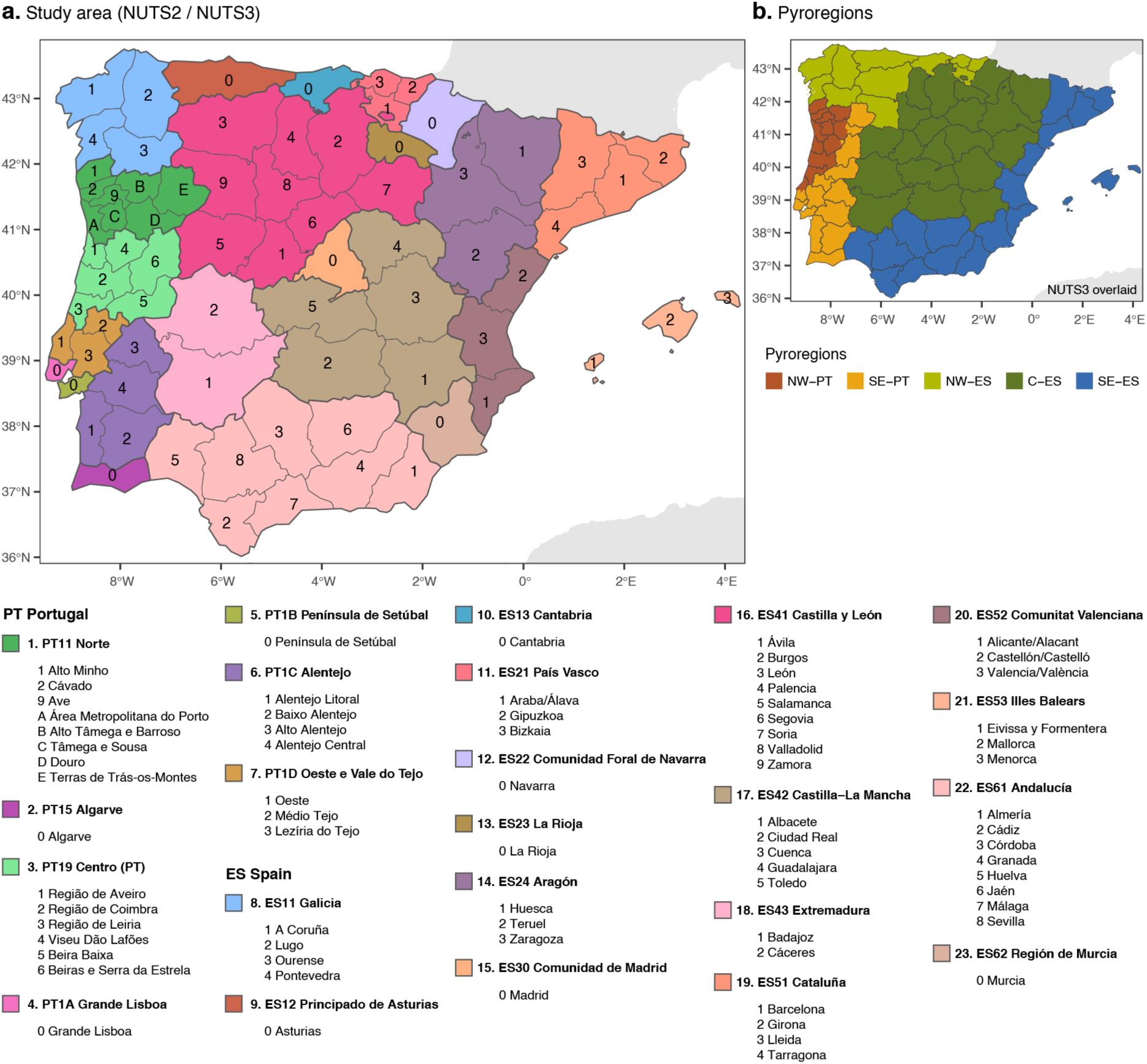
| Study area divided in NUTS2/NUTS3 administrative regions (a) and pyroregions (b).

**Figure S8.**
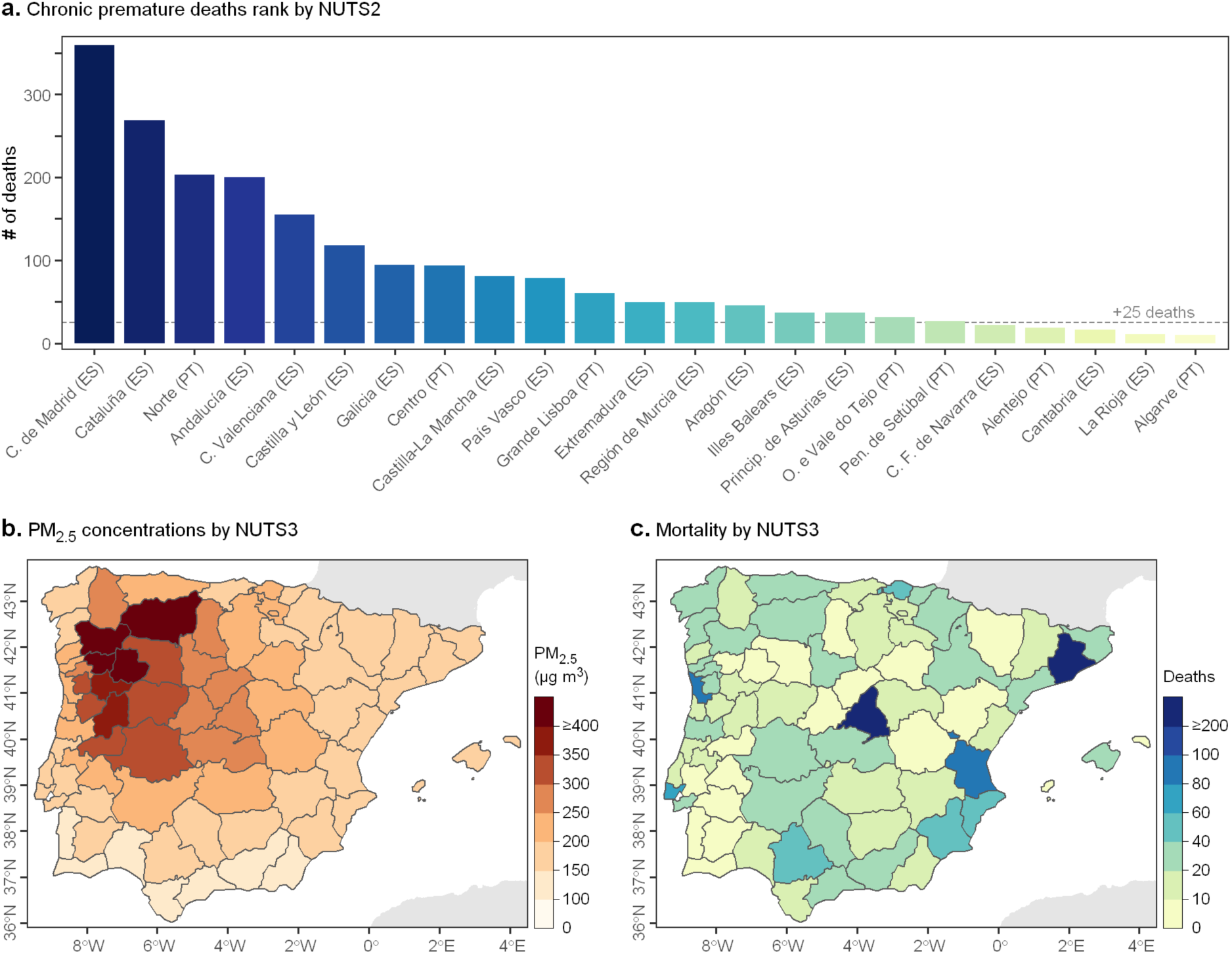
| Mortality associated with fire-derived PM2.5 across NUTS2 (a) and NUTS 3 (b and c) regions.

## Notes

### Competing Interest Statement

The authors have declared no competing interest.

